# A Facile Alternative Sustainable Process for the Extraction and Purification of Microbial Melanin

**DOI:** 10.1101/2021.05.10.443376

**Authors:** Vishal Ghadge, Pankaj Kumar, Tapan K. Maity, Kamalesh Prasad, Pramod B. Shinde

## Abstract

Conventional techniques for the extraction of the melanin encounter several disadvantages such as poor extraction efficiency, excessive use of acid and alkali, and lengthy time duration. To extract melanin in a more sustainable manner, herein suitability of an aqueous solution of quaternary ammonium hydroxide known as tetrabutylammonium hydroxide (40% w/w TBAOH in water) to extract and purify the pigment from the endophyte *Streptomyces hyderabadensis* 7VPT5-5R was investigated. The solvent could extract 5.54±0.03 g/L of pure melanin against 3.31± 0.02 g/L of pure melanin by conventional method from bacterial culture resulting 66% increase in the yield of the pigment. The quaternary ammonium based solvent was further recycled for 5 consecutive cycles and production of melanin with yield ranging from 5.54±0.03 g/L to 5.47± 0.02 g/L was obtained. The yield of the pigment was 38−40% higher in the recycled solvent in comparison to the melanin yield extracted by conventional method. Considering the efficiency of the solvent for effective extraction and purification of melanin and its recyclability potential, it can be considered as a sustainable solvent for the extraction of the pigments from bacterial culture. Furthermore, the developed method for extraction of melanin from microorganisms as a sustainable source would help in reducing the cost of melanin leading to its efficient utilization in diverse fields.

## INTRODUCTION

Melanins are dark black to brown coloured, hydrophobic, negatively charged, natural heterogeneous pigments which are constantly encountered in animals, plants, and in most of the microorganisms.^1^ The increased interest in melanin research is justified by its numerous biofunctions, such as colouration, camouflage, photo- and radio-protection, energy transducer, antioxidant, thermoregulator, chemoprotectant, antitumor, antiviral, antimicrobial, immunostimulant, anti-inflammatory, and detoxification.^1^ Microbial melanins have advantage over synthetic melanin and other natural melanins derived from plants and animals, in terms of easy and safe production, eco-friendliness, low cost, raw culture medium components, and biodegradablity.^2^

In the natural domain, three classes of melanin are frequently discerned by its colour and type of cyclic ring present. Eumelanins are black to brown in colour formed by oxidation of L-tyrosine (and/or L-phenylalanine) to 5,6-dihydroxyindole (DHI) or 5,6-dihydroxyindole-2-carboxylic acid (DHICA).^3^ Pheomelanins are yellow to red in colour and its synthesis is similar to eumelanins, wherein benzothiazine and benzothaizole units are formed from polymerization of cysteinyl DOPA (dihydroxyphenylalanine).^4^ Allomelanins, heterogenous group of polymers, are synthesized by oxidation/polymerization of 1,8-dihydroxynaphthalene (1,8-DHN) or 1,3,6,8-tetrahydroxynaphthalene (1,3,6,8-THN) via the pentaketide pathway.^5^

Melanin extraction is a challenging job on the grounds of its diversity and structural complexity. Generally employed base-dissolution and acid-precipitation method abide issues like low extraction efficiency, time consuming, multiple steps, use of corrosive solvents, and increased production cost. Also, the use of acidic- and basic-solution quite often leads to destruction of the molecule. Therefore, advanced extraction methods such as ultrasonic-assisted extraction (UAE), microwave-assisted extraction (MAE), and enzymatic digestion have been used to extract melanin.^6−8^However, these advanced methods are also associated with some drawbacks; for example, use of sequential digestion steps with multiple specific enzymes would prolong the producing time that ultimately raise the production cost of melanin; and rapid increase in temperature of the extraction mixture in microwave-assisted extraction method may terminate the extraction process early due to the boiling of the solvent.^9^ Even though, there are a number of reported extraction methods for different types of sources, the occurrence of melanin as bound form with cellular components and its insolubility in most of the organic solvent complicates extraction procedures. Thus, the desired compounds are not sufficiently diffused from the material into the solvent and consequently the extraction yield is reduced.^10^ The above insubstantial conventional production and extraction techniques result into limited melanin yield and subsequent high cost, which creates a barrier in its widespread industrial applicability. Therefore, there is an urgent need to develop a sustainable method to extract melanin from microbial source.

Although extraction of biomolecules directly using a quaternary ammonium electrolytes (QAEs) is not reported but solvents known as ionic liquids and deep eutectic solvents prepared using quaternary ammonium compounds are known to extract macromolecules such as carrageenan, agarose, and glaucarubinone.^11−13^ However, these solvents systems are emerging as a very good medium for the dissolution of biopolymers such as cellulose.^14,15^ The quaternary ammonium compounds being a hydrogen bond acceptor are known to react with hydrogen bond donors such as alcohols, acids, and sugars to form new structures by hydrogen bonding. In the previous study, we considered 40% aqueous solution of tetrabutylammonium hydroxide (40% w/w TBAOH in water) an ionic liquid but considering the fact that upon evaporation of water present in it, it was not possible to get a molten salt, hence herein, we are calling the solution as a quaternary ammonium base solution rather than ionic liquid.^16^ The quaternary ammonium base solution is expected to form favourable hydrogen bond with eumelanin due to the presence of carboxylic and hydroxyl groups in the structure and supposed for the latter’s selective extraction in the solvent (Figure S1). Certain imidazolium based ionic liquids are also reported to extract melanin from Alpaca fibre.^17^ Fungal polyketide pigment was also extracted by using quaternary and immidazolium based ionic liquid with aqueous two-phase system.^18^

In the present study, extracellular melanin was extracted from the culture broth of endophytic actinomycete *Streptomyces hyderabadensis* 7VPT5-5R using aqueous solution of TBAOH. Melanin production in the microbial culture was increased by optimizing medium components. The obtained melanin was characterized as eumelanin using physico-chemical and spectroscopic methods. The solvent was recycled for 5 consecutive cycles and yield of melanin in each of the cycles was monitored. The developed extraction method is simple, faster, one pot and eco-friendly, and has potential for large-scale extraction of pigments from natural resources.

## EXPERIMENTAL SECTION

### Chemicals and Materials

Tetrabutylammonium hydroxide [N(C_4_H_9_)_4_^+^ OH^−^] (40% w/w in H_2_O) was purchased from Molychem, Mumbai, India. Synthetic melanin was purchased from Sigma-Aldrich (St. Louis, MO, USA). Solvents were purchased from Qualigens (India). Microbial media components were purchased from HiMedia (India). All the chemicals used were of AR grade and were used as received without further purification.

### Isolation of endophytic actinomycetes

Plant *Salicornia brachiata* Roxb. was collected from Victor port, Amreli district, Gujarat on Feb. 26, 2018 (N 20° 58’ 53.2″, E 071° 33’ 21.2″). Collected plant sample was packed in sterile Ziploc bags and brought to laboratory in an ice-box for immediate processing as per reported method.^19,20^ Briefly, sample was cleaned and washed with water and Tween 20 (0.1% v/v) followed by surface sterilization sequentially with 4% NaOCl, 2.5% Na_2_S_2_O_3_, 70% ethyl alcohol, and sterile distilled water. The plant sample was subjected to continuous drying at 100 °C for 15 min and then plant parts were cut into small pieces and approximately 2 g of plant material was macerated in 1X PBS (pH 7). The plant extract was further treated with 1.5 % v/v phenol followed by an incubation of 15 min at 28 °C.^21^ Dilutions were prepared up to 10^−4^ right after this step and 100 µL of these dilutions were spread on ISP-4 agar supplemented with nystatin and cycloheximide (60 μg/mL each). The inoculated plates were incubated at 28 °C for 2−4 weeks, following which colonies appearing on ISP-4 agar plates were picked up.

### Microbial Culture for Production of Melanin

#### Screening for Melanin Production

Isolates were inoculated into ISP-6 broth composed of peptic digest of animal tissue 15 g, proteose peptone 5 g, yeast extract 1 g, ferric ammonium citrate 0.5 g, K_2_HPO_4_ 1 g, Na_2_S_2_O_3_ 0.08 g, pH 6.7±0.2 for 1 L and tyrosine broth (ISP-7) composed of glycerol 15 g, L-asparagine 1 g, L-tyrosine 0.5 g, K_2_HPO_4_ 0.5 g, MgSO_4_·7H_2_O 0.5 g, NaCl 0.5 g, and trace salts (FeSO_4_·7H_2_O 1.36 mg, CuCl_2_·2H_2_O 0.027 mg, CoCl_2_·6H_2_O 0.04 mg, Na_2_MoO_4_·2H_2_O 0.025 mg, ZnCl_2_ 0.02 mg, H_3_BO_3_ 2.85 mg, MnCl_2_·4H_2_O 1.8 mg, C_4_H_4_Na_2_O_6_ 1.77 mg), pH 7.3±0.1 for 1 L, incubated at 28 °C for 7−10 days. Production of brown to black pigment in the medium was scored as positive for melanin production. Melanin pigment production was quantified by measuring O.D. of the supernatant of culture broth spectrophotometrically at 400 nm.^22^

#### Identification of Melanin Producer

Melanin producing endophytic isolate was characterized by 16S rRNA analysis. The genomic DNA was isolated using Genetix DNA isolation kit. 16S rRNA gene was amplified using universal primers (27F 5’-AGAGTTTGATCCTGGCTCAG-3’, 1492R 5’GGTTACCTTGTTACGACTT-3’). PCR was performed using a thermal cycler (BioRad T100 Thermal Cycler) under the following optimized conditions: initial denaturation at 95 °C for 7 min, followed by 30 cycles of denaturation at 94 °C for 1 min, annealing at 57 °C for 1 min, and extension at 72 °C for 1 min, with a final extension temperature of 72 °C for 8 min. The 16S rRNA gene sequence was identified by a database search using the BLAST program (National Center for Biotechnology Information; http://www.ncbi.nlm.nih.gov).

#### Medium Optimization for Melanin Production

Medium components for melanin production were optimized by one factor at a time (OFAT) method. This method is used to identify the effect of individual component in which only one factor or a single variable is changed at a time while keeping all other variables constant.^23^ Total 23 medium components were screened one by one to check the effect of each component on melanin production. Tyrosine broth was used as basal medium containing glycerol as carbon source, both L-tyrosine and L-asparagine as nitrogen source. To identify the best carbon source, glycerol was replaced with six other carbon sources (sucrose, starch, lactose, fructose, maltose, and glucose). Similarly, the nitrogen source (L-tyrosine and L-asparagine) was replaced with eight other nitrogen sources (L-tyrosine, peptone bacteriological, yeast extract, ammonium sulphate, L-asparagine, casein acid hydrolysate, beef extract, potassium nitrate, sodium nitrate) to identify best nitrogen source. Trace salts of tyrosine broth were completely replaced with eight individual trace salts (ZnSO_4_, Co(NO_3_)_2_, CuSO_4_, MnCl_2_, FeSO_4_, NiCl_2_, CuCl_2_, and Ni(NO_3_)_2_). The effect of the medium components on melanin production was quantified by measuring the absorbance of the respective culture supernatants at 400 nm.^19^

The seed culture was prepared using the production medium composed of glycerol 20 g, NaNO_3_ 10 g, MgSO_4_ 1 g, K_2_HPO_4_ 1 g, NaCl 1 g, MnCl_2_ 0.004 g, pH 7.2. Then, 3 % (v/v) seed culture was transferred to 250 mL of production medium in a 1 L flask and the flask was incubated at 28 °C with shaking at 200 rpm for 8 days.

### Extraction of Melanin

After 8 days of incubation, culture broth was harvested, cell biomass was separated using centrifugation at 8000 rpm for 15 min, and the supernatant was collected. Then, the obtained supernatant was freeze dried to obtain crude powder using VirTis BenchTop K lyophilizer.

#### Conventional Method

In the conventional method, 1 mL water was added to 20 mg lyophilized powder obtained from the culture broth, the pH of the solution was adjusted to 12.0 with 1N NaOH and stirred for 1 h. Then, the pH of the solution was adjusted again to 2.0 using 6M HCl and allowed to stand for 4 h and then the solution was centrifuged at 13,000 rpm for 10 min. The obtained pellet was washed twice with ethanol followed by water to remove impurities. Finally, the purified melanin was obtained after lyophilizing the pellets collected in the previous step.

#### TBAOH-based Extraction

To 20 mg of lyophilized powder obtained from the culture broth, 1 mL water was added and mixed well followed by addition of 1 mL TBAOH and stirred for 30 min. To this mixture, 1 mL ethyl alcohol was added, mixed well, and then allowed to stand for 1 h followed by centrifugation at 13,000 rpm for 10 min. Then, the pellet was separated and washed twice with ethyl alcohol to remove impurities and the remaining supernatant was used for recycling of TBAOH. Eventually, the pellet was lyophilized to yield pure melanin.

#### Recycling of TBAOH

The supernatant containing TBAOH remained after precipitation of melanin was used for recycling of TBAOH. The equal amount of ethyl alcohol was added to supernatant facilitated easy evaporation of water using a rotary evaporator along with ethyl alcohol thus, leaving behind TBAOH only. Furthermore, efficiency and suitability of recycled TBAOH for the extraction of melanin was also investigated.^14^

### Characterization of Microbial Melanin

#### Melanin Biosynthesis

Kojic acid, a well-known melanin-synthesis inhibitor, was used to identify the enzyme involved in melanin biosynthesis. Concentrations of kojic acid ranging from 20−50 µg/mL were prepared in water and added to tyrosine broth. 1% (v/v) inoculum was added into above media and incubated at 28 °C, 160 rpm for 4−10 days and observed for pigment inhibition.^24^

#### Scanning Electron Microscopy (SEM) and Energy-Dispersive X-ray spectroscopy (EDX)

Extracted microbial melanin was dispersed in ethyl alcohol, sonicated for 5 min, and placed onto an SEM specimen stub. Then, the sample was dried in a desiccator overnight followed by visualization using JEOL JSM-7100F instrument employing 18 kV accelerating voltage.

#### Elemental Analysis

The percent elemental composition of carbon, hydrogen, nitrogen, and sulfur in the purified melanin was determined with 5 mg of melanin powder using an elemental analyzer (Elementar-Vario MICRO cube).^19^

#### UV/VIS Spectrophotometry

The extracted pure melanin 0.5 mg was dissolved in 1 mL of 0.5 N NaOH solution and the UV spectrum was recorded by scanning at wavelength range of 200−800 nm using 0.5 N NaOH solution as blank control using a Thermo Scientific Evolution 201 UV-Vis spectrophometer.^22^

#### Fourier Transform Infrared Spectroscopy (FT-IR)

The purified melanin was ground with KBr powder in an agate mortar to break it up and the mixed disc was scanned at 4000–400 cm^-1^ using a PerkinElmer FT-IR spectrometer (SpectrumGX, GSA).^25^

#### Electron Paramagnetic Resonance Spectroscopy (EPR)

The presence of free radical in purified melanin was analyzed by MS-5000 benchtop ESR/EPR spectrometer (Magnettech). 2 mg melanin powder was added into a thin-walled glass tube and EPR signals were recorded. The parameters used to acquire the spectra were as follows; modulation amplitude: 0.16 mT; modulation frequency: 100 kHz; center field: 325 mT; sweep width: 25 mT; sweep time: 2 min; microwave frequency: 9.1 GHz; microwave power: 0.1 mW; at room temperature.^26^

#### NMR Spectroscopy

10 mg of purified melanin was added into 1 mL of 40% sodium deuteroxide solution (Sigma-Aldrich) and sonicated for 2−3 min. The obtained solution was centrifuged at 13,000 rpm for 5 min and the dark brown coloured supernatant was used for NMR analysis.^27^ Bruker Avance II 500 MHz and Jeol 600 MHz NMR spectrometers were used to acquire ^1^H-NMR and ^13^C-NMR spectra, respectively.

#### Statistical Analysis

One-way ANOVA was conducted using IBM SPSS Statistic (version 27, 2004, IBM Inc., USA) Data are presented as means ± SE. When appropriate, significance of differences between means was determined using Tukey’s HSD. Differences of p < 0.05 were considered significant and p < 0.001 were considered as very significant.

## Results AND DISCUSSION

### Isolation of Endophytic Actinomycetes

The strain 7VPT5-5R was isolated on ISP-4 media from the halophyte *S. brachiata* Roxb. The pre-treatments helped to eliminate fast-growing gram-negative and other bacteria, allowing isolation of comparatively slow-growing actinomycetes on ISP-4 medium. After 7 days of incubation, the spore like colonies were picked up and purified on ISP-4 agar plate.

### Microbial Culture for Production of Melanin

#### Screening for Melanin Production

The isolate 7VPT5-5R produced dark brown colour after 4 days of incubation. The absorbance of culture supernatant measured by UV/VIS spectroscopy wherein higher absorbance was observed in the UV region, which was found to be decreased towards the visible region, which is the characteristic of melanin.^22^

#### Identification and Phylogenetic Analysis

The sequence of 16S rRNA (1364 bp) was amplified and sequenced at Macrogen (Seoul, South Korea). Based on BLAST result, endophytic isolate 7VPT5-5R was found to be belonged to genus Streptomyces and showed maximum sequence similarity with *S. hyderabadensis* OU-40 (99.55 %), *S. rubrogriseus* NBRC 15455 (98.72 %), and *S. rubrogriseus* DSM 41477 (98.72 %). Based on phylogenetic tree (Figure S2) analysis, the strain *S. hyderabadensis* 7VPT5-5R fall into a clade together with *S. hyderabadensis* OU-40 and formed a distinct phyletic line together with the strains *S. olivaceus* NBRC 12805, *S. pactum* NBRC 13433, and *S. litmocidini* NRRL B-3635.

#### Medium optimization for melanin production

Since culture media components influence the melanin production, screening of critical components are important for yield improvement.^25^ Among the six carbon sources, glycerol showed maximum melanin production (Figure S3). Similarly, the maximum melanin production was also observed in the culture containing sodium nitrate as nitrogen source. Among the trace elements, MnCl_2_ was found to show maximum melanin production. The ability to utilise various media components to achieve better melanin production varies from species to species. Various studies have reported that the yield of microbial melanin is significantly influenced by the nutrient components and culture conditions.^19,22,23,28^

### Extraction of melanin

After 8 days of incubation, culture broth was harvested and centrifuged at 8000 rpm for 15 min. The obtained culture supernatant was freeze dried to yield crude powder.

#### Conventional and TBAOH-based Extraction

As shown in Scheme 1, from 1 L culture broth, 12.511 g lyophilized crude powder was obtained from which 20 mg powder was individually used in the present study. The TBAOH-based extraction method resulted in a final quantity of 8.87 mg pure melanin from 20 mg lyophilized powder; thus, giving the melanin yield of 5.548± 0.03 g/L after 196 h incubation. In comparison to TBAOH-based extraction method, conventional method yielded less quantity (5.3 mg) of pure melanin from 20 mg lyophilized powder, thus, resulting the yield of 3.315± 0.02 g melanin from 1 L culture broth after 196 h incubation. There was an enhancement of 40% melanin content in the TBAOH-based extraction method. The reduced melanin yield in the conventional method can be asserted to the very low solubility of the extract containing melanin in water. Due to the insolubility of melanin, other associated proteins and components also get precipitated with melanin when conventional method is used. It is also noticed that the impurities in the melanin obtained using conventional extraction method are difficult to separate by washing with solvents. Furthermore, as shown in Figure S1, TBAOH acts as a hydrogen bond acceptor and forms preferential hydrogen bonds with the carboxyl- and hydroxyl-groups of eumelanin resulting in selective separation of the molecule in the solvent.

**Scheme 1:**
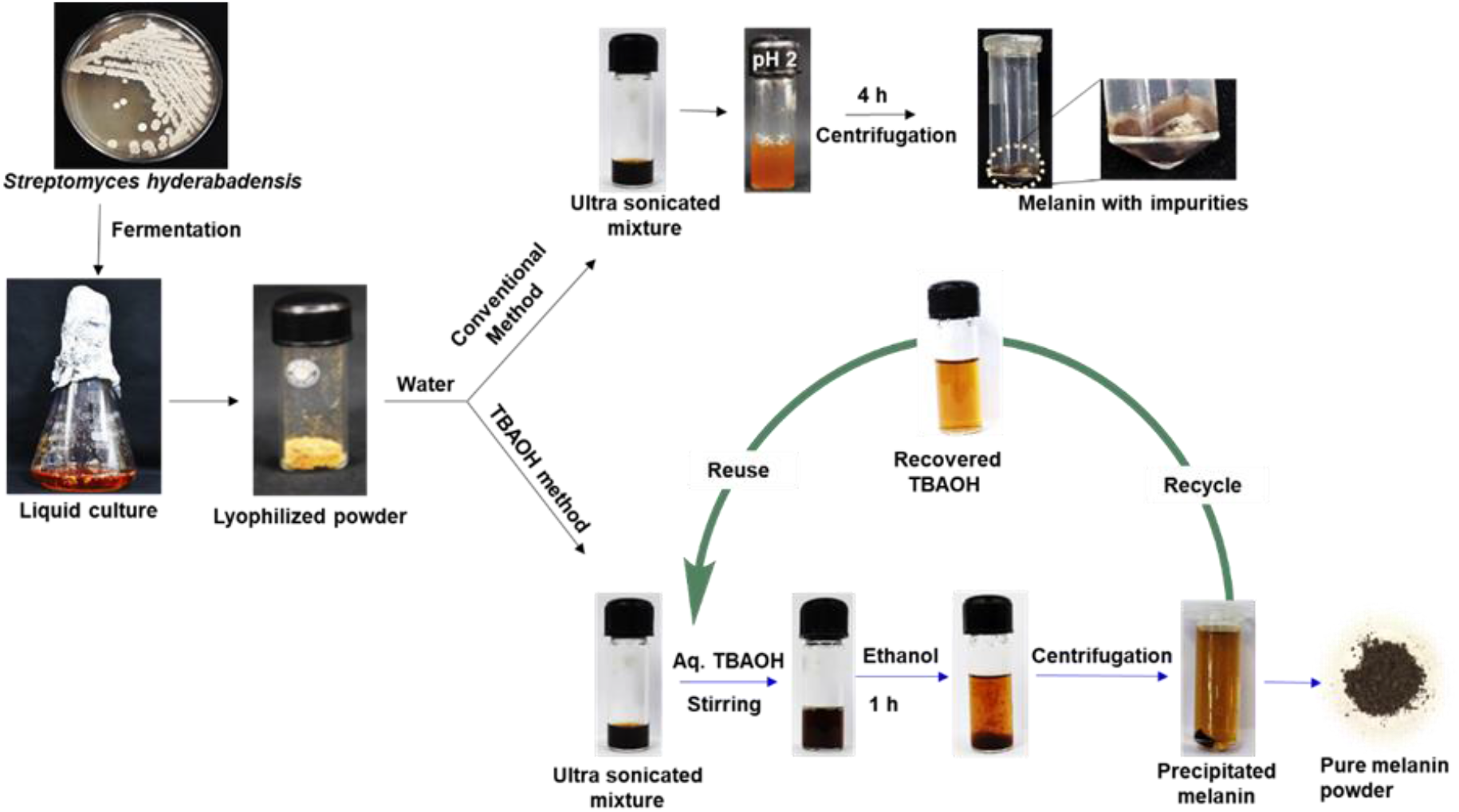
Extraction of melanin using conventional and TBAOH-based method.

#### Recycling of TBAOH

For recycling, the TBAOH remained after evaporation of ethyl alcohol and water was used for further dissolution of lyophilized crude powder obtained from the culture broth. The efficiency of recycled TBAOH was checked and its extraction yield was shown in Table 1. From the result, it was observed that the extraction efficiency of the recycled TBAOH is almost constant upto 5^th^ cycle of extraction. The above results fulfil the requirements of TBAOH for extraction of melanin and its extraction efficiency suggest that there is no significant loss of TBAOH during recycling. It is conceivable that this result will pave way for its industrial use for melanin extraction.

**Table 1.** Recycling of TBAOH and respective melanin yield.

| Numbers of recycles | Yield/ 20 mg of crude extract | Yield/L of culture |
| --- | --- | --- |
| 1 | 8.87 ± 0.03 mg | 5.54 ± 0.03 g |
| 2 | 8.83 ± 0.02 mg | 5.52 ± 0.02 g |
| 3 | 8.81 ± 0.02 mg | 5.51 ± 0.02 g |
| 4 | 8.78 ± 0.03 mg | 5.49 ± 0.03 g |
| 5 | 8.76 ± 0.03 mg | 5.47 ± 0.02 g |

#### Sustainability of Extraction Method towards Scale Up

In case of research on melanin, it is observed that the most of the work is focused on increasing its production yield from natural sources, but its extraction and purification is comparatively less explored. Among the various extraction methods reported for extraction of melanin, base-dissolution and acid-precipitation is the most commonly used method.^29,30^ Due to use of harsh acid treatment, melanin is found to be decomposed by extensive decarboxylation.^31^ Another detailed study employing spectroscopic analysis of the extracted melanin method concluded that base-dissolution and acid-precipitation method alters the molecular structure and attached metal coordination of extracted melanin.^32^ As a known fact, melanin is insoluble in most of the organic solvents and soluble at alkaline pH. Sometimes it was observed that due to high pH, melanin was found to be oxidized and this resulted in melanin degradation leading to structure alteration. Along with base-dissolution and acid-precipitation, some other treatments like dialysis, column chromatography, and washing with organic solvents are also reported for extraction and purification of melanin.^28^ Above methods have several disadvantages like time consuming, multistep, low extraction yield, and structural distortion. The advanced method likes cavitation-based extraction (CE), ultrasound-assisted extraction (UAE), and microwave-assisted extraction (MWE) were also employed for melanin extraction, but sonication heat was reported to be generated leading to decrease in the diffusion of melanin in the solvent resulting into reduced yield.^7,33,34^ Furthermore, for large scale extraction of melanin, the concentration of ultrasound waves cannot be kept constant. In conclusion, all above discussed methods are not facile for scale up of melanin extraction from natural sources. Herein, aqueous TBAOH-based method was optimized for selective extraction of melanin from microbial broth. The TBAOH solvent is recyclable, reusable and ecofriendly. Furthermore, all the experiments involving TBAOH for melanin extraction were carried out at room temperature. All above indications suggest suitability of this solvent (TBAOH) for industrial use for dissolution and extraction of melanin from biomass.

### Characterization of Microbial Melanin

#### Melanin Biosynthesis

The purpose of this study was to traverse the biosynthetic pathway of melanin using kojic acid, which is a DOPA melanin (eumelanin) pathway inhibitor. The eumelanin is synthesized by the enzyme tyrosinase.^23^ The inhibition of pigment production (Figure 1) was observed after treatment with kojic acid in the culture medium indicating that the melanin synthesized by endophyte *S. hyderabadensis* 7VPT5-5R is eumelanin melanin.

**Figure 1.**
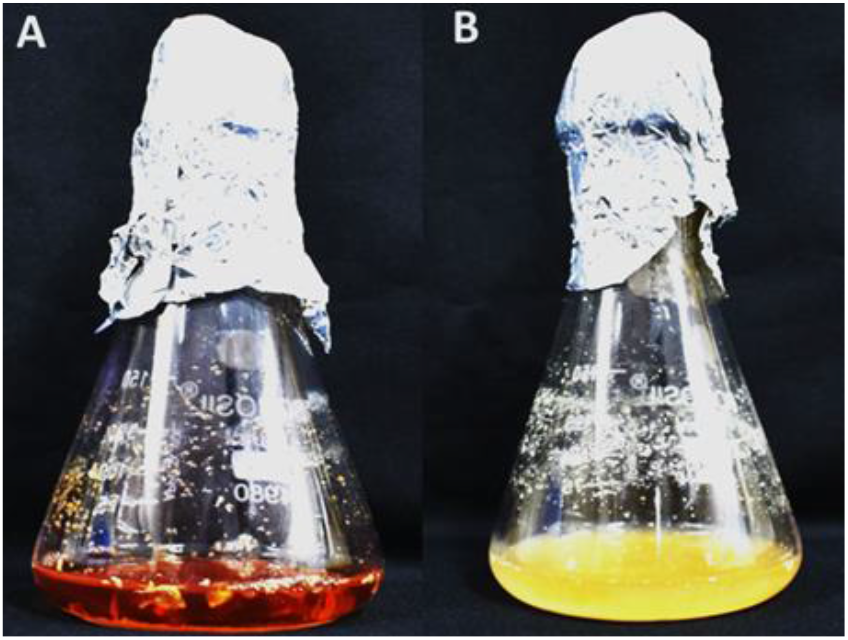
Melanin synthesis inhibition: Control (A) and Treatment with kojic acid (B).

#### Scanning Electron Microscopy (SEM) and Energy-Dispersive X-ray spectroscopy (EDX)

In the SEM study, purified melanin showed an amorphous (irregular) shape (Figure 2) pattern in comparison with reported bacterial purified melanin.^19^ The EDX technique is helpful to determine the composition of samples based on the peak emission of X-rays resulting from the interaction between each element of a compound and the electron beam.^35^ The major elemental composition of the melanin surface revealed by EDX analysis includes carbon, nitrogen, oxygen, and sulfur with approximate weight as 32.95%, 5.10%, 50%, and 0.1%, respectively.

**Figure 2.**
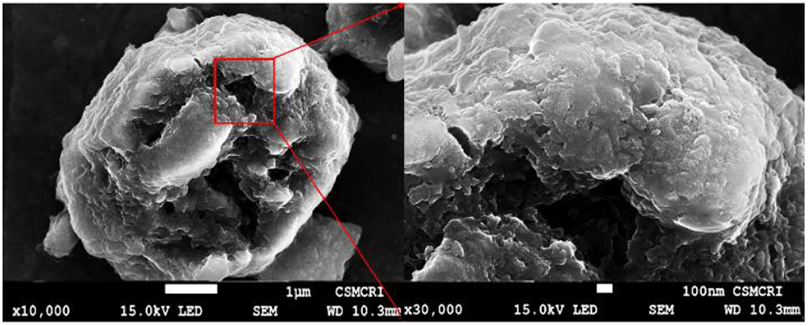
SEM image of purified melanin at different magnification.

#### Elemental Analysis

Elemental analysis of the isolated melanin showed percent composition of C, H, N, and S as 27.38, 4.50, 5.08, and 0.88%, respectively. The presence of large proportion of C, H, N and the absence of S, indicated that the obtained melanin from endophytic actinomycete *S. hyderabadensis* 7VPT5-5R can be classified as eumelanin.^19^

#### UV/VIS Spectrophotometry

The maximum absorbance was observed at 229 nm, which was decreased towards the visible region (Figure 3) which is the characteristic property of melanin. The *λ*max of microbial melanin varies from species to species in the range of UV region.^22^ The ratio of *A*_650_/*A*_500_ helps to calculate the amount of eumelanin in the total melanin.^19^ The ratio *A*_650_/*A*_500_ was calculated to be 0.35 and indicated that extracted melanin belong to eumelanin class, whereas, a ratio below 0.15 is reported for pheomelanin.

**Figure 3.**
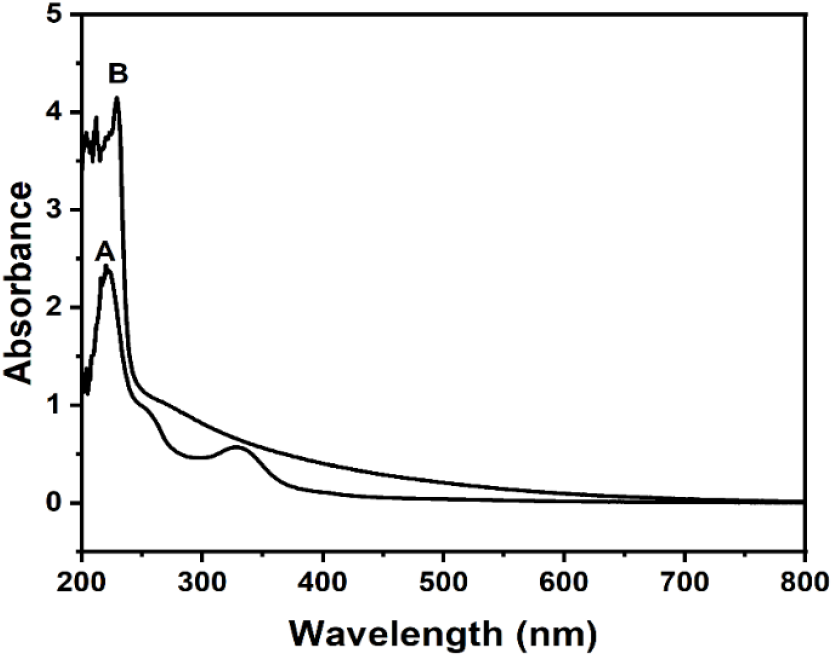
UV-Vis spectra of purified melanin (A) and synthetic melanin (B)

#### Fourier Transform Infrared Spectroscopy (FT-IR)

FT-IR is mainly used to identify functional groups to aid structure elucidation of melanin. The FT-IR spectrum (Figure 4) of purified melanin exhibited a broad band at 3422.23 cm^−1^ due to the stretching vibrations of O-H and N-H groups, a small band at 2964.23 cm^−1^ due to stretching vibration of aliphatic C-H groups.^22^ The characteristic bands appeared at 1661.74 cm^−1^ corresponds to the C=O stretching or aromatic C=C stretching.^36^ The peaks at 1452.56 cm^−1^ and 1405.90 cm^−1^ are attributed to a C-N stretching, indicating that it has the typical indole structure of melanin.^24^ The peak at 1042.76 cm^−1^ indicated the presence of aliphatic CH group in plane, which is characteristic of melanin pigment.^22^ The presence of aromatic C-H group corresponds to band at 877.13 cm^−1^. The band appeared at 707.62 cm^−1^ corresponds to alkene C-H substitution in the melanin pigment.^29^

**Figure 4.**
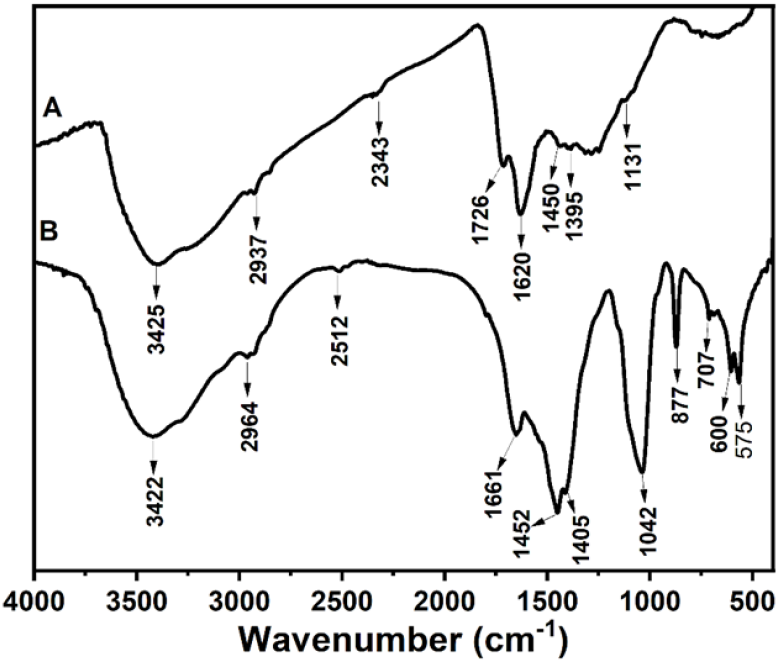
FT-IR spectra of synthetic melanin (A) and purified melanin (B)

#### Electron Paramagnetic Resonance Spectroscopy (EPR)

EPR spectrum revealed that obtained melanin particles contained stable free radicals, which is characteristic physiological property. The EPR spectrum (Figure 5) of the purified melanin showed the g-factor and the signal peak was observed at 2.003 and 337 mT, respectively, indicating the presence of free radicals. Furthermore, EPR spectroscopy has been successfully used to identify the type of melanin on the basis of g value and EPR line shapes.^37^ In this study, the obtained melanin was identified as eumelanin by comparison of EPR line shapes to the line shapes obtained from the synthetic eumelanin (Figure 5).

**Figure 5.**
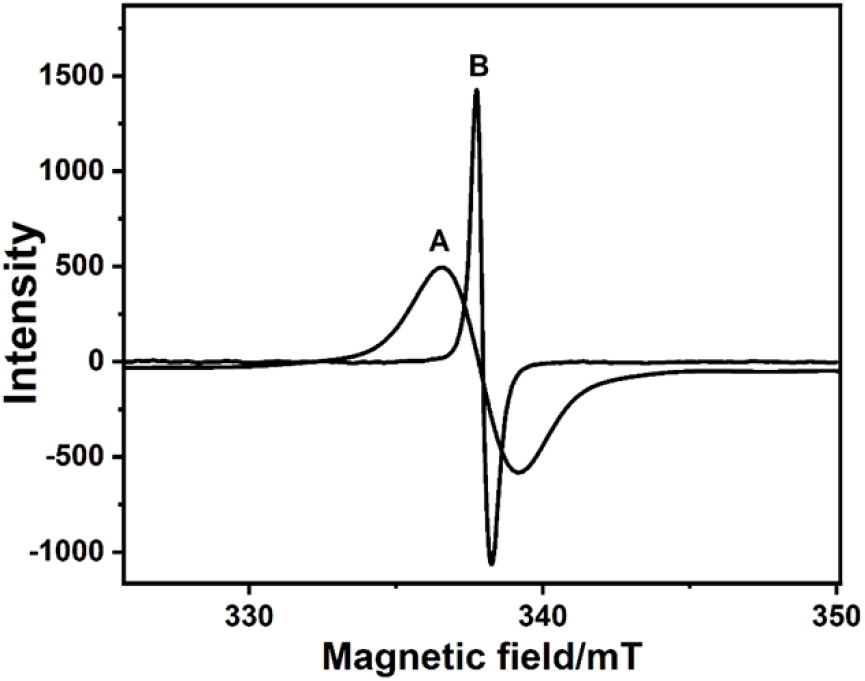
EPR spectra of purified melanin (A) and synthetic melanin (B).

#### NMR Spectroscopy

The ^1^H NMR spectrum (Figure 6) of isolated melanin from *S. hyderabadensis* 7VPT5-5R was found to display signals for broad functional groups. Thus, the peaks observed at δ_H_ 0.5−1.5 were assigned to CH_3_ and CH_2_ groups of alkyl fragments. Peaks from δ_H_ 3.16−4.13 were assigned to protons on carbons attached to oxygen or nitrogen atoms. In the aromatic region, the peaks at δ_H_ 6.52−6.75 were assigned to the benzene-CH group of indole. The peak observed at δ_H_ 7.20−7.81 indicated presence of heterocyclic ring and complexity of melanin.^38^ The single signal observed at δ_H_ 8.46 indicated the existence of pyrrole-CH group of a carboxyl substituted indole.^39^ For comparison, the ^1^H NMR spectrum of the synthetic melanin is shown in Figure S4.

**Figure 6.**
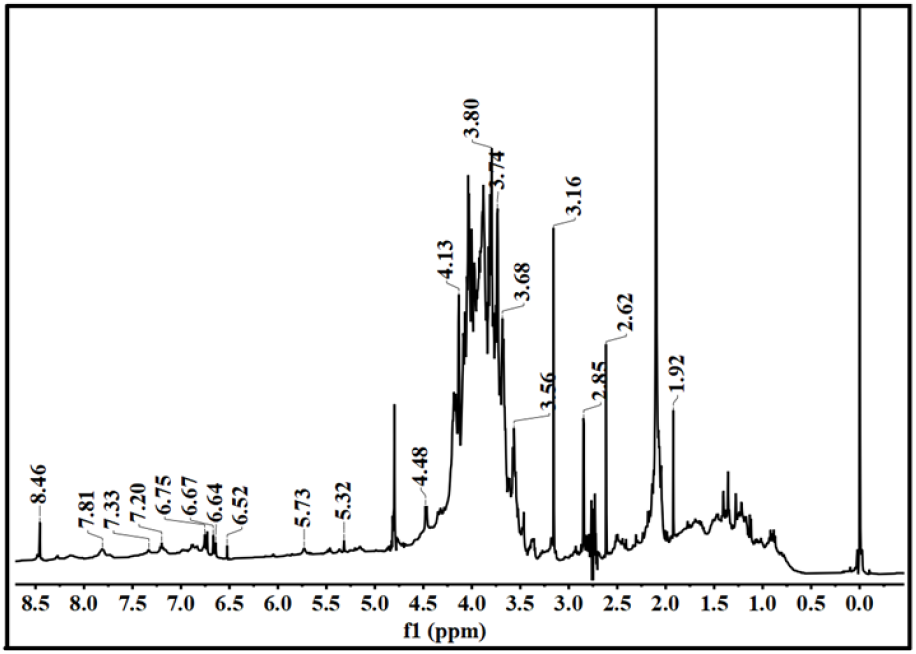
^1^H NMR spectrum of purified melanin.

## CONCLUSIONS

In conclusion, a facile alternative sustainable process for the extraction of microbial melanin from culture broth of the endophytic actinomycete was developed. The melanin-producing endophytic actinomycete was isolated from the halophyte *Salicornia brachiata* Roxb. and subsequently identified as *Streptomyces hyderabadensis* 7VPT5-5R. The production yield of melanin was optimized using OFAT method and the extracted melanin was identified as eumelanin on the basis of UV/VIS, FT-IR, EPR, NMR, and elemental analysis. An aqueous solution of TBAOH was investigated for extraction of melanin and it was further shown that the TBAOH can be recycled for reuse in extraction of melanin. It was also observed that the yield of the extracted melanin was 38−40% higher in the recycled solvent comparison to conventional method. The aqueous TBAOH solvent is recyclable, reusable, and eco-friendly suggesting its suitability for industrial use for dissolution and extraction of melanin from biomass.

## ASSOCIATED CONTENT

### Supporting Information

Addition experimental details, interaction of aqueous TBAOH and melanin, phylogentic analysis of *S. hyderabadensis* 7VPT5-5R, media component effects on melanin production, and ^1^H NMR spectrum of synthetic melanin.

## AUTHOR INFORMATION

### Notes

There are no conflicts to declare.

## ACKNOWLEDGEMENTS

This work was supported by Scientific and Engineering Research Board (SERB), Department of Science and Technology [ECRA/2016/000788 and EEQ/2016/000268]; and Council of Scientific and Industrial Research (CSIR), New Delhi [MLP/0027]. VAG thanks SERB-DST for a project fellowship (GAP-2050 and GAP-2057). PK acknowledges DBT-JRF fellowship from Department of Biotechnology (GAP-2074). TKM Thanks CSIR for a project fellowship (HCP0019). Analytical and Environmental Science Division and Centralized Instrument Facility of the Institute is acknowledged for providing instrumentation facilities.

## TOC graphic

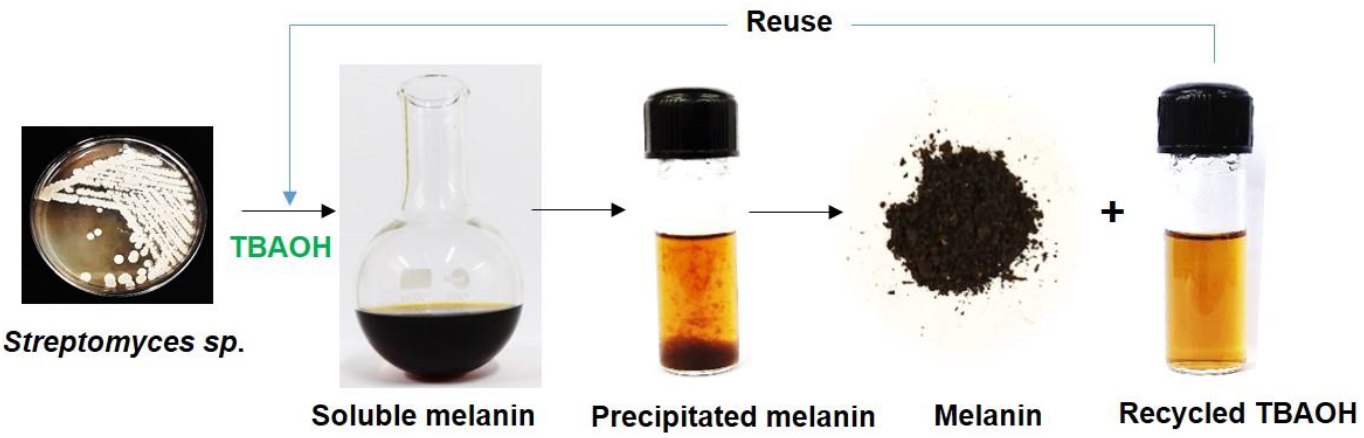

## REFERENCES

(1) Singh, S.; Nimse, S. B.; Mathew, D.; Dhimmar, A.; Sahastrabudhe, H.; Gajjar, A.; Ghadge, V.; Kumar P.; Shinde, P. B. Microbial melanin: Recent advances in biosynthesis, extraction, characterization, and applications. Biotechnol. Adv. 2021, 107773. In Press. DOI: 10.1016/j.biotechadv.2021.107773

(2) Sen, T.; Barrow, C. J.; Deshmukh, S. K. Microbial pigments in the food industry−challenges and the way forward. Front. Nutr. 2019, 6, 7. DOI: 10.3389/fnut.2019.00007

(3) Xie, W.; Pakdel, E.; Liang, Y.; Kim, Y. J.; Liu, D.; Sun, L.; Wang, X. Natural eumelanin and its derivatives as multifunctional materials for bioinspired applications: A review. Biomacromolecules 2019, 20, 4312–4331. DOI: 10.1021/acs.biomac.9b01413

(4) Ito, S.; Kolbe, L.; Weets, G.; Wakamatsu, K. Visible light accelerates the ultraviolet A-induced degradation of eumelanin and pheomelanin. Pigment Cell Melanoma Res. 2019, 32, 441–447. DOI: 10.1111/pcmr.12754

(5) Varga, M.; Berkesi, O.; Darula, Z.; May, N. V.; Palagyi, A. Structural characterization of allomelanin from black oat. Phytochem. 2016, 130, 313–320. DOI: 10.1016/j.phytochem.2016.07.002

(6) Zou, Y.; Xie, C.; Fan, G.; Gu, Z.; Han, Y. Optimization of ultrasound-assisted extraction of melanin from Auricularia auricula fruit bodies. Innov. Food Sci. Emerg. Technol. 2010, 11, 611–615. DOI: 10.1016/j.ifset.2010.07.002

(7) Lu, Y.; Ye, M.; Song, S.; Li, L.; Shaikh, F.; Li, J. Isolation, purification, and anti-aging activity of melanin from Lachnum singerianum. Biotechnol. Appl. Biochem. 2014, 174, 762–771. DOI: 10.1007/s12010-014-1110-0

(8) Punitha, V.; Kannan, P.; Saravanabhaven, S.; Thanikaivelan, P.; Rao, J.; Saravanan, P.; Nair, B. Enzymatic removal of melanin in enzyme based dehairing and fiber opening. J. Am. Leather Chem. Assoc. 2008, 103, 203–208.

(9) Chuyen, H. V.; Nguyen, M. H.; Roach, P. D.; Golding, J. B.; Parks, S. E. Microwave-assisted extraction and ultrasound-assisted extraction for recovering carotenoids from Gac peel and their effects on antioxidant capacity of the extracts. Nutr. Food Sci. 2018, 6, 189–196. DOI: 10.1002/fsn3.546

(10) Nguyen, V. T.; Vuong, Q. V.; Bowyer, M. C.; Van Altena, I. A.; Scarlett, C. J. Microwave-assisted extraction for saponins and antioxidant capacity from Xao tam phan (Paramignya trimera) root. J. Food Process. Preserv. 2017, 41, e12851. DOI: 10.1111/jfpp.12851

(11) Das, A. K.; Sequeira, R. A.; Maity, T. K.; Prasad, K. Bio-ionic liquid promoted selective coagulation of κ-carrageenan from Kappaphycus alvarezii extract. Food Hydrocoll. 2021, 111, 106382. DOI: 10.1016/j.foodhyd.2020.106382

(12) Sharma, M.; Chaudhary, J. P.; Mondal, D.; Meena, R.; Prasad, K. A green and sustainable approach to utilize bio-ionic liquids for the selective precipitation of high purity agarose from an agarophyte extract. Green Chem. 2015, 17, 2867–2873. DOI: 10.1039/c4gc02498b

(13) Kholiya, F.; Bhatt, N.; Rathod, M. R.; Meena, R.; Prasad, K. Fundamental studies on the feasibility of deep eutectic solvents for the selective partition of glaucarubinone present in the roots of Simarouba glauca. J. Sep. Sci. 2015, 38, 3170–3175. DOI: 10.1002/jssc.201500470

(14) Zhong, C.; Wang, C.; Huang, F.; Jia, H.; Wei, P. Wheat straw cellulose dissolution and isolation by tetra-n-butylammonium hydroxide. Carbohydr. Polym. 2013, 94, 38–45. DOI: 10.1016/j.carbpol.2013.01.043

(15) Zhang, J.; Wu, J.; Yu, J.; Zhang, X.; He, J.; Zhang, J. Application of ionic liquids for dissolving cellulose and fabricating cellulose-based materials: state of the art and future trends. Mater. Chem. Front. 2017, 1, 1273–1290 DOI: 10.1039/C6QM00348F

(16) Singh, N.; Prasad, K. Multi-tasking hydrated ionic liquids as sustainable media for the processing of waste human hair: a biorefinery approach. Green Chem. 2019, 21, 3328–3333. DOI: 10.1039/C9GC00542K

(17) Liang, Y.; Han, Q.; Byrne, N.; Sun, L.; Wang, X. Recyclable one-step extraction and characterization of intact melanin from alpaca fibers. Fibers Polym. 2018, 19, 1640–1646. DOI: 10.1007/s12221-018-8144-9

(18) Lebeau, J.; Venkatachalam, M.; Fouillaud, M.; Dufossé, L.; Caro, Y. Extraction of fungal polyketide pigments using ionic liquids. In Book of Abstracts, 2016 8^th^ International Conference of Pigments in Food, Coloured foods for health benefits., Cluj-Napoca, Romania.

(19) Ghadge, V.; Kumar, P.; Singh, S.; Mathew, D. E.; Bhattacharya, S.; Nimse, S. B.; Shinde, P. B. Natural melanin produced by the endophytic Bacillus subtilis 4NP-BL associated with the halophyte Salicornia brachiata. J. Agric. Food Chem. 2020, 68, 6854–6863. DOI: 10.1021/acs.jafc.0c01997

(20) Singh, S.; Ghadge, V. A.; Kumar, P.; Mathew, D. E.; Dhimmar, A.; Sahastrabudhe, H.; Nalli, Y.; Rathod, M. R.; Shinde, P. B. Biodiversity and antimicrobial potential of bacterial endophytes from halophyte Salicornia brachiata. Anton. Leeuw. Int. J. G. 2021, 114, 591–608. DOI: 10.1007/s10482-021-01544-4

(21) Hayakawa, M.; Sadakata, T.; Kajiura, T.; Nonomura, H. New methods for the highly selective isolation of Micromonospora and Microbispora from soil. J. Ferment. Bioeng. 1991, 72, 320–326. DOI: 10.1016/0922-338X(91)90080-Z

(22) El-Naggar, N. E. A.; El-Ewasy, S. M. Bioproduction, characterization, anticancer and antioxidant activities of extracellular melanin pigment produced by newly isolated microbial cell factories Streptomyces glaucescens NEAE-H. Sci. Rep. 2017, 7, 1–19. DOI: 10.1038/srep42129

(23) Wang, L.; Li, Y.; Li, Y. Metal ions driven production, characterization and bioactivity of extracellular melanin from Streptomyces sp. ZL-24. Int. J. Biol. Macromol. 2019, 123, 521–530. DOI: 10.1016/j.ijbiomac.2018.11.061

(24) Manivasagan, P.; Venkatesan, J.; Senthilkumar, K.; Sivakumar, K.; Kim, S. K. Isolation and characterization of biologically active melanin from Actinoalloteichus sp. MA-32. Int. J. Biol. Macromol. 2013, 58, 263–274. DOI: 10.1016/j.ijbiomac.2013.04.041

(25) Al Khatib, M.; Harir, M.; Costa, J.; Baratto, M. C.; Schiavo, I.; Trabalzini, L.; Pollini, S.; Rossolini, G. M.; Basosi, R.; Pogni, R. Spectroscopic characterization of natural melanin from a Streptomyces cyaneofuscatus strain and comparison with melanin enzymatically synthesized by tyrosinase and laccase. Molecules 2018, 23, 1916. DOI: 10.3390/molecules23081916

(26) Suwannarach, N.; Kumla, J.; Watanabe, B.; Matsui, K.; Lumyong, S. Characterization of melanin and optimal conditions for pigment production by an endophytic fungus, Spissiomyces endophytica SDBR-CMU319. PLoS One 2019, 14, e0222187. DOI: 10.1371/journal.pone.0222187

(27) Katritzky, A. R.; Akhmedov, N. G.; Denisenko, S. N.; Denisko, O. V. ^1^H NMR spectroscopic characterization of solutions of Sepia melanin, Sepia melanin free acid and human hair melanin. Pigment Cell Res. 2002, 15, 93–97. DOI: 10.1034/j.1600-0749.2002.1o062.x

(28) Madhusudhan, D. N.; Mazhari, B. B. Z.; Dastager, S. G.; Agsar, D. Production and cytotoxicity of extracellular insoluble and droplets of soluble melanin by Streptomyces lusitanus DMZ-3. BioMed Res. Int. 2014, 2014, Article ID 306895. DOI: 10.1155/2014/306895

(29) Sajjan, S.; Kulkarni, G.; Yaligara, V.; Lee, K.; Karegoudar, T. B. Purification and physiochemical characterization of melanin pigment from Klebsiella sp. GSK. J. Microbiol. Biotechnol. 2010, 20, 1513–1520. DOI: 10.4014/jmb.1002.02006

(30) Sun, S.; Zhang, X.; Sun, S.; Zhang, L.; Shan, S.; Zhu, H. Production of natural melanin by Auricularia auricula and study on its molecular structure. Food Chem. 2016, 190, 801–807. DOI: 10.1016/j.foodchem.2015.06.042

(31) Ito, S. Reexamination of the structure of eumelanin. Biochim Biophys Acta Gen Subj. 1986, 883, 155–161. DOI: 10.1016/0304-4165(86)90146-7

(32) Liu, Y.; Kempf, V. R.; Brian Nofsinger, J.; Weinert, E. E.; Rudnicki, M.; Wakamatsu, K.; Ito, S.; Simon, J. D. Comparison of the structural and physical properties of human hair eumelanin following enzymatic or acid/base extraction. Pigment Cell Res. 2003, 16, 355–365. DOI: 10.1034/j.1600-0749.2003.00059.x

(33) Zou, Y.; Xie, C.; Fan, G.; Gu, Z.; Han, Y. Optimization of ultrasound-assisted extraction of melanin from Auricularia auricula fruit bodies. Innov. Food Sci. Emerg. Technol. 2010, 11, 611–615. DOI: 10.1016/j.ifset.2010.07.002

(34) Lu, M.; Yu, M.; Shi, T.; Ma, J.; Fu, X.; Meng, X.; Shi, L. Optimization of ultrasound-assisted extraction of melanin and its hypoglycemic activities from Sporisorium reilianum. J. Food Process. Preserv. 2020, 44, e14707. DOI: 10.1111/jfpp.14707

(35) Correa, N.; Covarrubias, C.; Rodas, P. I.; Hermosilla, G.; Olate, V. R.; Valdés, C.; Meyer, W.; Magne, F.; Tapia, C. V. Differential antifungal activity of human and cryptococcal melanins with structural discrepancies. Front. Microbiol. 2017, 8, 1292. DOI: 10.3389/fmicb.2017.01292

(36) Hou, R.; Liu, X.; Xiang, K.; Chen, L.; Wu, X.; Lin, W.; Zheng, M.; Fu, J. Characterization of the physicochemical properties and extraction optimization of natural melanin from Inonotus hispidus mushroom. Food chem. 2019, 277, 533–542. DOI: 10.1016/j.foodchem.2018.11.002

(37) Zdybel, M.; Pilawa, B.; Drewnowska, J. M.; Swiecicka, I. Comparative EPR studies of free radicals in melanin synthesized by Bacillus weihenstephanensis soil strains. Chem. Phys. Lett. 2017, 679, 185–192. DOI: 10.1016/j.cplett.2017.05.013

(38) El-Gamal, M. S.; El-Bialy, H. A.; Elsayed, M. A.; Khalifa, M. A. Isolation and characterization of melanized yeast form of Aureobasidium pullulans and physiological studies on the melanization process. J. Nucl. Sci. Technol. 2017, 5, 57–72.

(39) Saini, A. S.; Tripathi, A.; Melo, J. S. On-column enzymatic synthesis of melanin nanoparticles using cryogenic poly (AAM-co-AGE) monolith and its free radical scavenging and electro-catalytic properties. RSC Adv. 2015, 5, 47671–47680. DOI: 10.1039/C5RA18965A

